# Kinase-specific phosphorylation of tau outside the amyloid core encodes fibril fold selection

**DOI:** 10.64898/2026.08.04.742669

**Authors:** Reshma Ramesh, Sambhasan Banerjee, Madhu Babu Gajula Balija, Pijush Chakraborty, Christian Dienemann, Meytal Landau, Markus Zweckstetter

**Affiliations:** German Center for Neurodegenerative Diseases (DZNE), Von-Siebold-Str. 3a, 37075 Göttingen, Germany; CSSB Centre for Structural Systems Biology, Deutsches Elektronen-Synchrotron DESY, Hamburg, Germany; Department of Biotechnology and Bioinformatics, University of Hyderabad, Hyderabad, 500046, Telangana, India; Department of Molecular Biology, Max Planck Institute for Multidisciplinary Sciences, Am Faßberg 11, 37077 Göttingen, Germany; Department of Biology, Technion-Israel Institute of Technology, Haifa, Israel; The Center for Experimental Medicine, Universitätsklinikum Hamburg-Eppendorf (UKE), Hamburg, Germany; Department for NMR-based Structural Biology, Max Planck Institute for Multidisciplinary Sciences, Am Faßberg 11, 37077 Göttingen, Germany; Department of Neurology, University Medical Center Göttingen, University of Göttingen, Waldweg 33, 37073, Göttingen, Germany

## Abstract

Tauopathies are defined by the accumulation of filamentous tau assemblies that adopt disease-specific molecular conformations, or tau strains. Although tau is extensively hyperphosphorylated in Alzheimer’s disease, how phosphorylation patterns influence tau folding and strain selection remains unclear. Here, we show that site-specific phosphorylation of 0N3R tau by extracellular signal-regulated kinase 2 (ERK2), occurring predominantly within the proline-rich region and largely excluding the microtubule-binding domain, is sufficient to direct tau assembly into a structurally homogeneous fibril conformation. Residue-resolved NMR spectroscopy identifies a defined and quantifiable ERK2 phosphorylation pattern in the proline-rich region of tau. Cryo-electron microscopy at 3.0 Å resolution reveals that ERK2-phosphorylated 3R tau assembles into filaments with an ordered cross-β fold adopting an Alzheimer’s disease PHF fold, despite the absence of phospho-sites within the fibril core. These filaments exhibit robust seeding activity in tau biosensor cells. Together, these results demonstrate that kinase-specific phosphorylation outside the amyloid core can be sufficient to bias tau toward a defined fibril structure, establishing a direct mechanistic link between kinase specificity, post-translational modification, and tau strain formation.

## Introduction

Tauopathies are characterized by the accumulation of filamentous tau assemblies that adopt disease-specific molecular conformations^1,2^. High-resolution cryo-electron microscopy has revealed that tau filaments isolated from different tauopathies adopt distinct folds, implicating fibril structure as a determinant of pathological phenotype^2,3^. However, the molecular mechanisms that bias tau toward particular fibrillar architectures remain poorly understood.

Tau is an intrinsically disordered microtubule-associated protein that undergoes extensive post-translational modification, with phosphorylation being the most prominent in disease^4^. In Alzheimer’s disease and related tauopathies, phosphorylation is strongly enriched in the proline-rich region (PRR), whereas the microtubule-binding domain (MTBD), which forms the ordered cross-β core of tau fibrils, is comparatively sparsely modified^5–7^. While phosphorylation is known to promote tau aggregation, how phosphorylation patterns localized outside the amyloid core influence fibril fold selection remains unresolved.

One emerging concept is that post-translational modifications may influence tau aggregation by biasing early assembly steps rather than stabilizing the final fibril structure^8–10^. The proline-rich region is well positioned to mediate such effects. It is highly dynamic, engages in transient long-range interactions with the repeat domain, and regulates access to aggregation-prone sequence elements^11–13^. Modifications within this region could therefore influence the trajectory of tau assembly without becoming incorporated into the mature fibril core. However, direct experimental evidence linking defined kinase-specific phosphorylation patterns to tau fold selection has remained limited.

Extracellular signal-regulated kinase 2 (ERK2) is a proline-directed kinase implicated in neuronal signaling and tau pathology^14–16^. ERK2 preferentially phosphorylates tau at specific sites within the proline-rich region while exhibiting minimal activity toward the microtubule-binding domain, providing a tractable system to examine the structural consequences of non-core phosphorylation in isolation^17^.

Here, we combine residue-resolved NMR spectroscopy, quantitative phosphorylation analysis, and high-resolution cryo-electron microscopy to define the structural outcomes of ERK2-mediated tau phosphorylation. We show that ERK2 installs a specific phosphorylation pattern confined largely to the proline-rich region that markedly accelerates aggregation of 0N3R tau. Despite the absence of strong phosphorylation within the microtubule-binding domain, ERK2-phosphorylated 3R tau assembles into homogeneous filaments whose atomic structure closely resembles Alzheimer’s disease–associated tau fibrils and exhibits potent seeding activity in tau biosensor cells. These findings provide evidence that phosphorylation outside the amyloid core can be sufficient to bias tau toward a defined fibril fold, implicating kinase-specific non-core modification as a determinant of tau conformational diversity.

## Results

### Site-specific phosphorylation of 0N3R tau by human ERK2 kinase

To define the site-specific phosphorylation pattern generated by ERK2, recombinant human 0N3R tau was phosphorylated in vitro using purified human ERK2 kinase under conditions optimized to favor kinase-selective modification while avoiding extensive hyperphosphorylation. The overall extent of phosphorylation was assessed by SDS–PAGE (Fig. S1a), and intact mass spectrometry revealed two liquid chromatography (LC)-resolved populations of ERK2-phosphorylated tau, with the predominant species containing approximately five to six phosphate groups per molecule (Fig. S1 d,e).

Phosphorylation sites were initially identified by LC–MS/MS analysis of phosphorylated tau bands excised from SDS–PAGE. ERK2 predominantly targeted residues within the proline-rich region, including T111, T153, T175, T181 (AT270 epitope), S184, S191, S202, T212, T217, T231 (AT180 epitope), S235, and T245 (Fig. 1a). Additional phosphorylation was detected at several sites in the C-terminal region, including S396/S404 (PHF-1 epitope), S400, T403, and S422. Phosphorylation at T205, previously reported for ERK2^10^, was not detected in our dataset, likely due to the low peptide counts. Overall, the identified phosphorylation pattern was consistent with prior reports of ERK2 activity toward tau ^10,17^.

**Figure 1.**
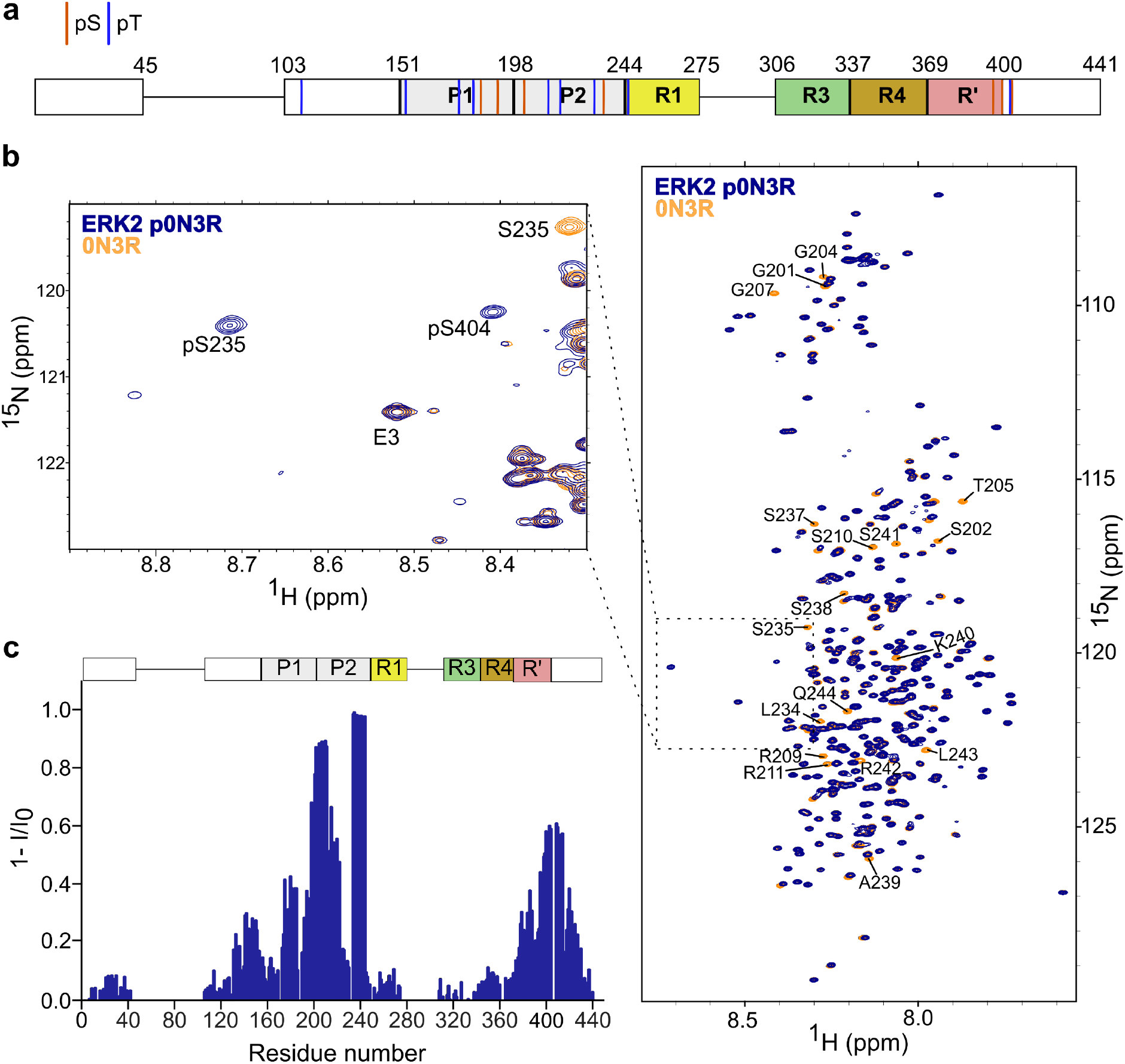
ERK2-specific phosphorylation pattern of 0N3R tau. **a,** Schematic domain diagram of human 0N3R tau with ERK2 phosphorylation sites identified by LC–MS/MS. Residue numbering follows the longest tau isoform 2N4R that additionally has two N-terminal inserts (residues 45-102) and the pseudo-repeat R3 (residues 275-305). Phosphorylated serine and threonine residues are indicated. **b,** Overlay of two-dimensional ^1^H–^15^N HSQC spectra of unmodified 0N3R tau (orange) and ERK2-phosphorylated 0N3R tau (blue), showing the appearance of new cross-peaks corresponding to phosphorylated residues and intensity changes at affected sites (right). Expanded region of the HSQC spectra highlighting representative phosphorylation-induced spectral changes (left). **c,** Residue-specific intensity attenuation (1 − I/I₀) derived from HSQC spectra, where I and I₀ denote peak intensities of the ERK2 phosphorylated and unmodified 0N3R tau, respectively, providing a quantitative measure of phosphorylation site occupancy and local effects.

Because mass spectrometry provides limited accuracy for quantifying residue-specific phosphorylation stoichiometry, we next employed NMR spectroscopy to determine the predominant phosphorylation sites and their relative occupancies. Two-dimensional ^1^H–^15^N HSQC spectra of ERK2-phosphorylated 0N3R tau exhibited pronounced spectral changes relative to unmodified tau, including the appearance of new cross-peaks characteristic of phosphorylated serine and threonine residues and marked signal attenuation at modified sites and neighboring residues (Fig. 1b). Residue-specific intensity changes, quantified as 1 − I/I₀, revealed a phosphorylation pattern closely matching the MS data (Fig. 1c).

Quantitative analysis showed that ERK2 efficiently phosphorylated multiple residues within the proline-rich region, including S235 (∼99%), S241 (∼98%), S210 (∼89%), T212 (∼86%), and the AT8 epitope residues S202 (∼86%) and T205 (∼88%). Residues spanning S235 to Q244 displayed particularly strong signal attenuation. For residues S237 and S238, reduced signal intensity was observed. However, direct phosphorylation at these sites could not be unambiguously assigned, as neither MS nor NMR data could distinguish direct modification from indirect effects of neighboring phosphorylation.

In contrast, phosphorylation within the C-terminal region occurred at substantially lower levels. Among these sites, S400 and S404 were phosphorylated to approximately 55% and 54%, respectively, whereas phosphorylation of S396 could not be directly quantified due to spectral overlap with T231 but was estimated to be ∼30% based on attenuation of adjacent resonances. Notably, phosphorylation levels at T231 and T181 were lower than reported in previous studies using ERK2 from Xenopus laevis^17^, likely reflecting both species-specific kinase differences and the reduced kinase concentrations used here to preserve site selectivity.

In summary, these analyses demonstrate that ERK2 generates a highly defined phosphorylation pattern on 0N3R tau that is strongly enriched in the proline-rich region, with only partial modification of the C-terminal domain and minimal involvement of the microtubule-binding repeats.

### ERK2 phosphorylation accelerates 3R tau aggregation and promotes twisted fibril morphology

Following site-specific characterisation of ERK2-mediated phosphorylation on 0N3R tau, we next examined its effect on 3R tau fibrillization using a previously established cofactor-free in vitro aggregation assay^18^. Aggregation reactions were performed in triplicate, and fibril formation was monitored by Thioflavin T (ThT) fluorescence.

ERK2 phosphorylation markedly accelerated 3R tau aggregation, as reflected by reduced aggregation half-times relative to unmodified tau (Fig. 2a,b). In contrast, pelleting assays performed at the end of the aggregation reactions showed comparable extents of fibrillization for ERK2-phosphorylated and unmodified 0N3R tau, with approximately 45% and 54 % of the protein recovered in the insoluble fraction, respectively. These results indicate that ERK2 phosphorylation primarily affects the rate and pathway of tau assembly rather than the final thermodynamic extent of fibril formation.

**Figure 2.**
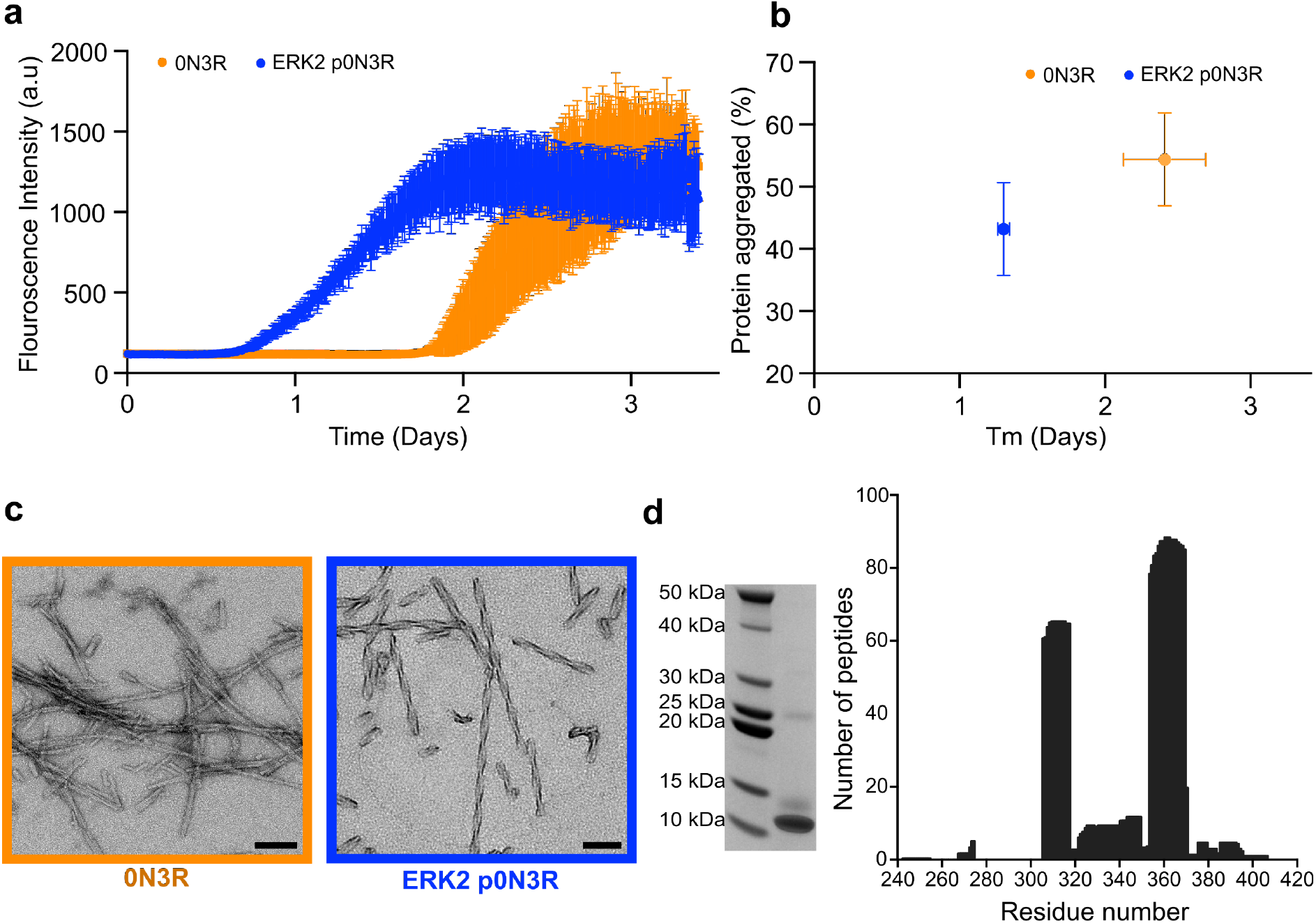
ERK2 phosphorylation accelerates tau aggregation and promotes twisted fibril morphology. **a,** Thioflavin T fluorescence traces showing aggregation kinetics of 25 μM ERK2-phosphorylated and unmodified 0N3R tau under cofactor-free conditions. Data represent mean ± s.d. from three independent experiments. **b,** Fraction of tau converted into insoluble aggregates plotted against aggregation half-time (Tₘ) for ERK2-phosphorylated and unmodified 0N3R tau. Aggregated fractions were determined by BCA analysis of supernatants following fibril pelleting. **c,** Representative negative-stain transmission electron microscopy images of fibrils formed by ERK2-phosphorylated and unmodified 0N3R tau. Scale bar, 100 nm. **d,** Analysis of the protease-resistant fibril core of ERK2-phosphorylated 0N3R tau. Left, SDS–PAGE of pronase-digested fibrils showing a prominent protease-resistant band. Right, number of peptides identified by mass spectrometry mapping to residues within the protease-resistant core. Values represent the mean from three independent experiments.

Negative-stain transmission electron microscopy revealed pronounced morphological differences between fibrils formed by phosphorylated and unmodified tau. ERK2-phosphorylated 0N3R tau predominantly assembled into twisted filaments, whereas unmodified tau primarily formed straight filaments (Fig. 2c). The twisted morphology observed for ERK2-phosphorylated tau is reminiscent of paired helical filament–like architectures reported for Alzheimer’s disease tau, although definitive structural comparison requires higher-resolution analysis.

To assess whether the twisted fibrils formed by ERK2-phosphorylated 0N3R tau exhibit a defined protease-resistant core, fibril samples were subjected to digestion with pronase, a mixture of broad-specificity proteases that preferentially removes solvent-accessible flexible regions while preserving the structured fibril core. SDS-PAGE analysis of the digestion products revealed a prominent protease-resistant band at approximately 12 kDa (Fig. 2d), consistent with the presence of an ordered fibril core.

Mass spectrometric analysis of the excised protease-resistant band showed that the core of ERK2-phosphorylated 0N3R tau fibrils predominantly comprises peptides spanning residues ∼306 to ∼400 (Fig. 2d), a region consistent with the core identified in tau filaments isolated from Alzheimer’s disease brain. A smaller number of peptides extending toward both the N-and C-terminal regions were also detected, likely reflecting limited accessibility of residues proximal to the core to proteolytic cleavage. Importantly, three independent aggregation experiments yielded highly similar twisted fibril morphologies and protease-resistant peptide profiles, underscoring the reproducibility of the fibril architecture formed by ERK2-phosphorylated tau.

These results show that ERK2-mediated phosphorylation accelerates tau aggregation and promotes the formation of a reproducible twisted fibril morphology with a defined protease-resistant core, supporting the existence of a constrained fibril fold that is further resolved by high-resolution structural analysis.

### ERK2-phosphorylated 0N3R tau assembles into PHF-like filaments with an Alzheimer’s disease fold

To determine the atomic structure of ERK2-phosphorylated 0N3R tau fibrils, we used cryo-electron microscopy (cryo-EM) and helical reconstruction (Table S1). Cryo-EM micrographs revealed abundant twisted fibrils embedded in thin vitreous ice (Fig. 3a). Three-dimensional reconstructions showed that ERK2-phosphorylated 0N3R tau fibrils are composed of two protofilaments arranged into a closed, C-shaped fold characteristic of Alzheimer’s disease paired helical filaments (PHFs) (Fig. 3b-d). The final cryo-EM reconstruction was resolved to an overall resolution of 3.02 Å. No alternative fibril morphologies were observed among the particles selected for reconstruction, indicating a high degree of structural homogeneity within the reconstructed population.

**Figure 3.**
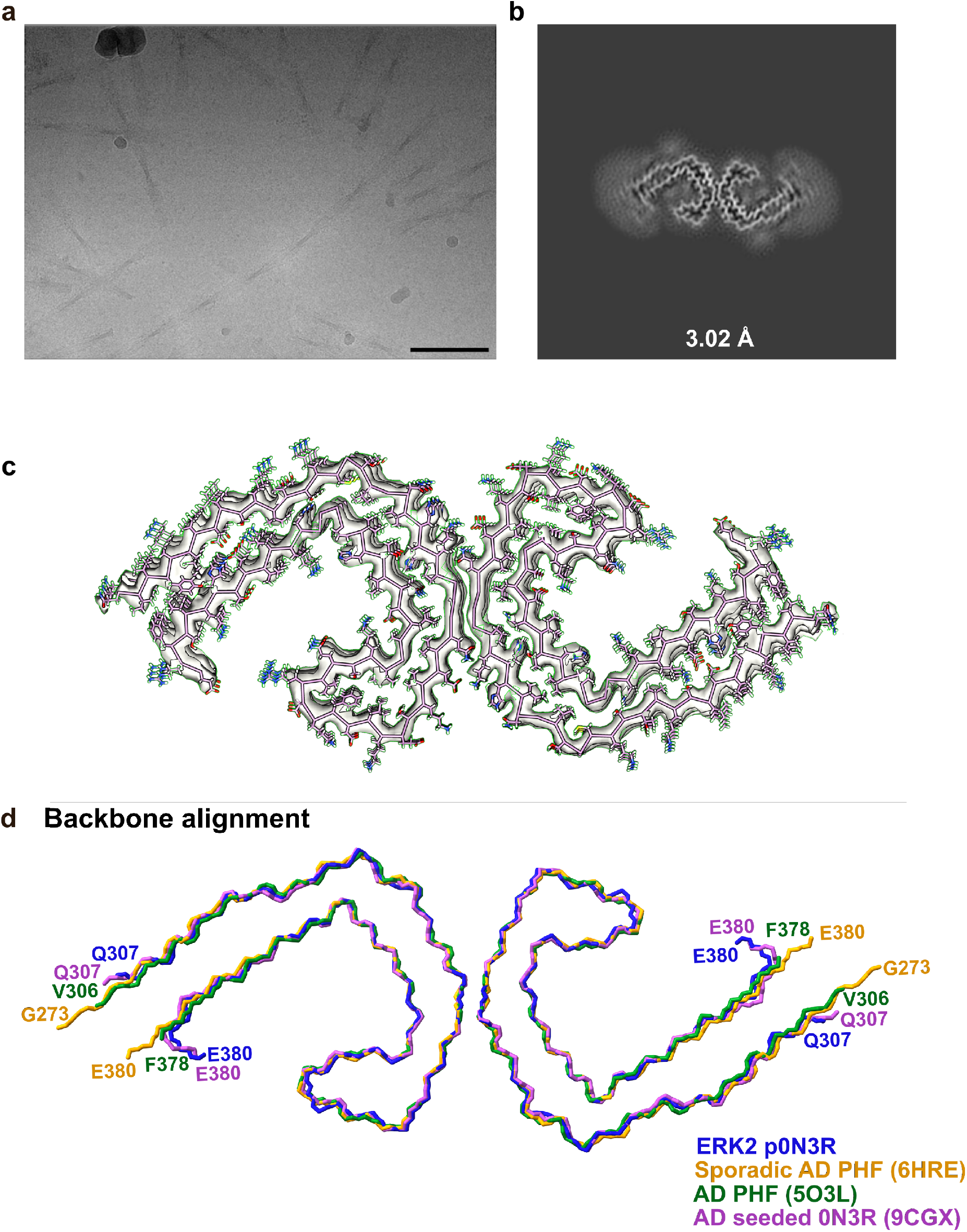
Cryo-EM structure of ERK2-phosphorylated 0N3R tau fibrils. **a,** Representative cryo-electron micrograph of ERK2-phosphorylated 0N3R tau fibrils. Scale bar, 500 Å. **b,** Central slice through the three-dimensional cryo-EM density map of ERK2-phosphorylated 0N3R tau fibrils, reconstructed to an overall resolution of 3.02 Å. **c,** Atomic model of ERK2-phosphorylated 0N3R tau fitted into the cryo-EM density map. **d,** Backbone superposition of ERK2-phosphorylated 0N3R tau fibrils with Alzheimer’s disease-derived paired helical filament (PHF) structures (PDB: 6HRE and 5O3L) and an Alzheimer’s disease-seeded 0N3R tau fibril structure (PDB: 9CGX), illustrating a high degree of structural similarity among the fibril folds.

Cross-sectional views of the reconstruction showed that the two protofilaments associate with a helical twist of approximately -0.52° and a rise of 2.37 Å along the fibril axis (Fig. S2a). Within each protofilament, β-strands are related by a helical rise of approximately 4.88 Å (Fig. S2b). ERK2-phosphorylated 0N3R tau fibrils exhibited an average cross-over distance of 860 ± 73 Å and a width of 161 ± 19 Å at the widest point of the helical repeat (Fig. S2a).

Assignment of the cryo-EM density to the tau sequence revealed a fibril core spanning residues Q307-E380. This core closely overlaps with those reported for PHFs isolated from Alzheimer’s disease brain, including the AD PHF structure encompassing residues V306–F378 (PDB: 5O3L)^19^ and the sporadic AD PHF structure spanning residues G304–E380 (PDB: 6HRE)^20^ (Fig. 3b-d). Backbone alignment of ERK2-phosphorylated 0N3R tau fibrils with these AD-derived PHF structures showed a high degree of structural similarity, with backbone RMSDs of 1.17 Å and 2.04 Å relative to 5O3L and 6HRE, respectively (Fig. 3d).

The primary structural deviation relative to AD brain–derived PHFs was observed at the C-terminal residues R379 and E380, which adopt an upward-oriented conformation in the ERK2-phosphorylated 0N3R structure. This difference may reflect electrostatic effects involving residues proximal to the fibril core boundary. When residues R379 and E380 were excluded from the alignment, the backbone RMSD relative to 6HRE decreased to 1.14 Å. Notably, the ERK2-phosphorylated 0N3R tau structure was nearly indistinguishable from the recently reported Alzheimer’s disease–seeded 0N3R tau fibril structure (PDB: 9CGX)^21^, with a backbone RMSD of 0.91 Å, which also exhibits a similar orientation of the C-terminal residues.

The data demonstrate that ERK2-mediated phosphorylation of 0N3R tau gives rise to a homogeneous fibril structure that adopts an Alzheimer’s disease PHF fold at near-atomic resolution.

### ERK2-phosphorylated 0N3R tau fibrils exhibit robust seeding activity

Tauopathies, including AD, propagate transneuronally via a “prion-like” mechanism, in which misfolded tau assembled in affected neurons can spread to healthy neurons and induce aggregation of endogenous tau^22,23^. To assess the seeding efficiency of ERK2-phosphorylated 0N3R tau fibrils, we used fluorescence resonance energy transfer (FRET)–based tau biosensor assay in HEK293T cells stably expressing CFP- and YFP-tagged tau repeat domains carrying the P301S mutation^24^. Seed-induced tau aggregation was monitored by FRET arising from intermolecular proximity of CFP- and YFP-tagged tau upon fibril-mediated nucleation (Fig. 4a).

**Figure 4.**
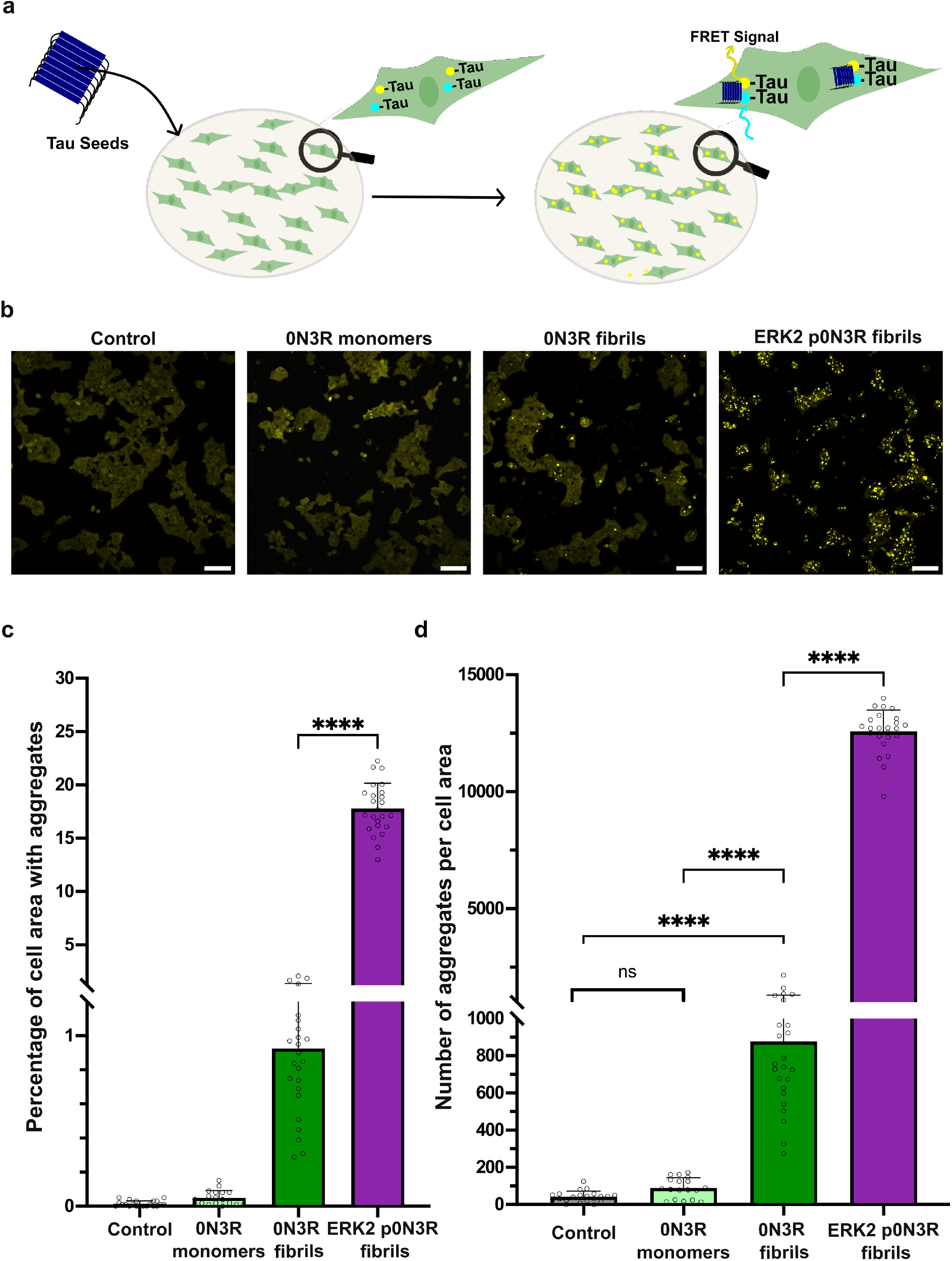
Seeding activity of ERK2-phosphorylated 0N3R tau fibrils in biosensor cell assay. **a,** Schematic of the FRET-based tau biosensor assay in HEK293T cells expressing CFP- and YFP-tagged 4R tau repeat domains, illustrating seed-induced aggregation and FRET signal generation. **b,** Representative FRET images of biosensor cells 24 h after transfection with ERK2-phosphorylated 0N3R tau fibrils, unmodified 0N3R tau fibrils, unmodified 0N3R tau monomers, and control (no transfection). Scale bar, 100 μm. All conditions were transfected with an equal total amount of tau (5 μg). **c,** Quantification of seeding efficiency based on the percentage of cell area containing FRET-positive aggregates. Data represent mean ± s.d. from three independent experiments. Statistical significance was assessed by one-way ANOVA (****P < 0.0001). **d,** Quantification of seeding efficiency based on the number of FRET-positive aggregates per unit cell area. Data represent mean ± s.d. from three independent experiments. Statistical significance was assessed by one-way ANOVA (****P < 0.0001).

Biosensor cells were transfected with ERK2-phosphorylated 0N3R tau fibrils, unmodified 0N3R tau fibrils, or unmodified 0N3R tau monomers, using an equal total amount of tau (5 µg) in each condition. Cells treated with ERK2-phosphorylated 0N3R tau fibrils exhibited pronounced FRET-positive aggregates 24 h after transfection, whereas substantially fewer aggregates were observed in cells treated with unmodified fibrils, and no detectable aggregation was observed in cells treated with monomeric tau or in untreated controls (Fig. 4b).

Quantitative analysis showed that ERK2-phosphorylated 0N3R tau fibrils produced a significantly higher fraction of FRET-positive cell area and a greater number of aggregates per unit area compared with unmodified 0N3R tau fibrils (Fig. 4c,d). These differences were reproducible across three independent experiments and were statistically significant by one-way ANOVA (P < 0.0001).

These results demonstrate that ERK2-phosphorylated 0N3R tau fibrils exhibit enhanced seeding efficiency in a cellular tau biosensor system compared with unmodified 0N3R tau fibrils, providing functional support for the distinct aggregation behavior of the structurally defined fibril state.

## Discussion

In this study, we define how kinase-specific phosphorylation outside the amyloid core can encode tau fibril fold selection. By combining residue-resolved NMR mapping of ERK2-mediated phosphorylation with near-atomic cryo-electron microscopy, we show that phosphorylation concentrated in the proline-rich region, with limited contribution from the C-terminal domain and minimal modification of the repeat region, is sufficient to bias 0N3R tau toward assembly into a single, dominant fibril structure adopting an Alzheimer’s disease paired helical filament (PHF) fold. These findings provide structural evidence that post-translational modification can act upstream of amyloid core formation to constrain tau conformational outcomes.

Previous studies have shown that phosphorylation modulates tau aggregation kinetics, oligomerization pathways, and strain-like behavior, but phosphorylation has typically been examined as a bulk modification state or via phosphomimetic substitutions, with structural consequences inferred indirectly from aggregation assays or low-resolution morphology^8,25–27^. As a result, it has remained unclear whether defined kinase-specific phosphorylation patterns alone are sufficient to specify a particular fibril fold. Our results address this gap by linking a quantitatively defined phosphorylation pattern introduced by a single kinase to a unique fibril structure resolved at near-atomic resolution.

A central insight from this work is that phosphorylation need not be incorporated into the ordered cross-β core to exert decisive control over fibril structure. Despite the near-complete absence of phospho-sites within the resolved amyloid core, ERK2-phosphorylated tau reproducibly assembles into a PHF fold that is nearly indistinguishable from tau filaments isolated from Alzheimer’s disease brain and from Alzheimer’s disease–seeded 0N3R tau fibrils. This argues against models in which phosphorylation stabilizes fibril folds through direct participation in the amyloid spine and instead supports a mechanism in which phosphorylation biases early assembly or nucleation events that determine the final fold^28,29^.

The proline-rich region is well positioned to mediate such upstream control. This region is highly dynamic, engages in transient long-range interactions with the repeat domain, and regulates access to aggregation-prone motifs that form the fibril core^30–32^. ERK2 installs a defined and quantitatively dominant phosphorylation pattern within this region while leaving the microtubule-binding repeats largely unmodified. The highly reproducible emergence of a single fibril fold from this defined modification state suggests that ERK2-mediated phosphorylation channels tau assembly along a restricted pathway, selecting a specific β-strand registry and protofilament interface (Fig. 5). Consistent with this interpretation, ERK2-phosphorylated tau fibrils exhibit minimal polymorphism across independent preparations and datasets, demonstrating that kinase-specific modification can suppress structural heterogeneity rather than merely accelerate aggregation.

**Figure 5.**
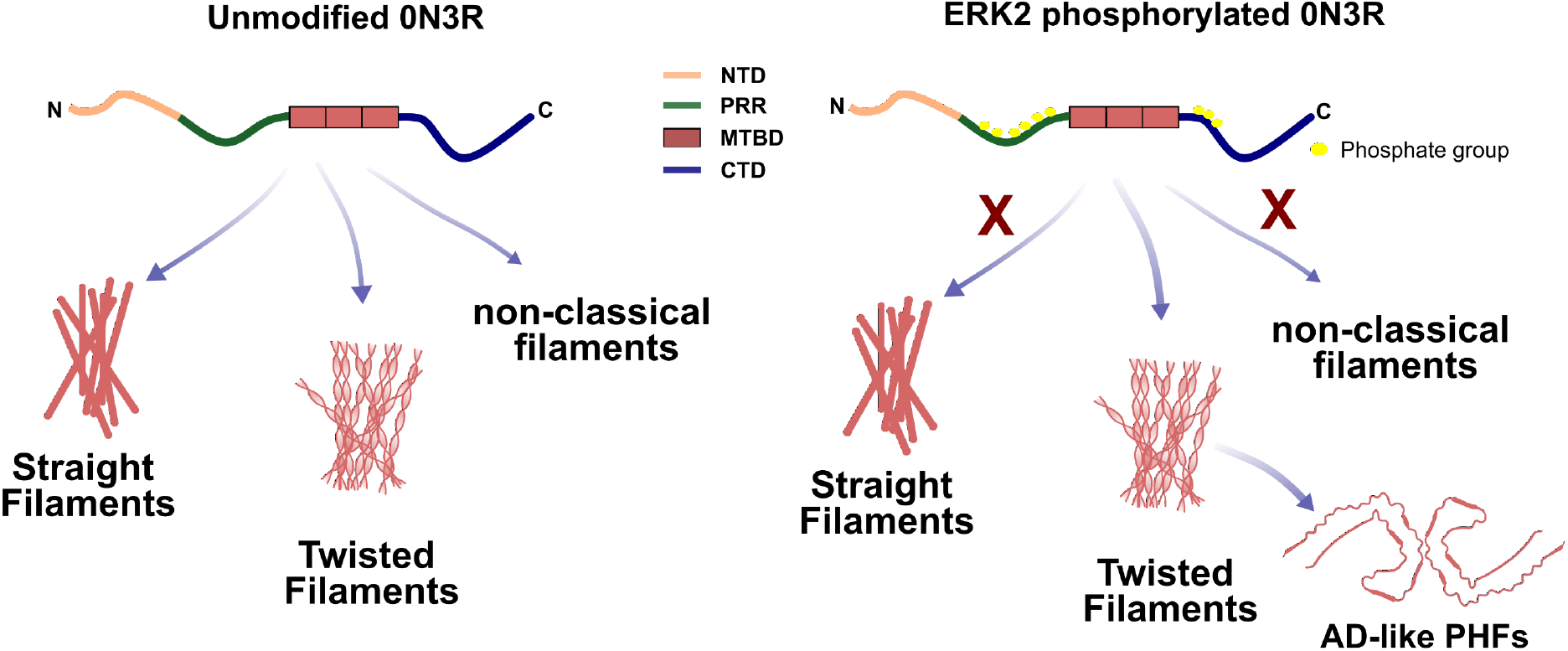
Kinase-specific phosphorylation outside the amyloid core encodes tau fibril fold selection. In the absence of defined post-translational modifications, intrinsically disordered tau samples multiple aggregation pathways, leading to structurally heterogeneous fibril polymorphs. ERK2-mediated phosphorylation establishes a modification pattern concentrated within the proline-rich region (PRR), with little phosphorylation of the C-terminal domain and minimal phosphorylation of the microtubule-binding repeat domain that forms the amyloid core. This spatially restricted phosphorylation biases early assembly events and narrows the spectrum of accessible aggregation pathways. As a result, ERK2-phosphorylated 0N3R tau reproducibly assembles into a single, highly homogeneous twisted fibril that adopts an Alzheimer’s disease PHF-like fold, despite the absence of phosphosites within the ordered cross-β core

Recent work has independently shown that kinase-mediated phosphorylation can bias tau toward PHF-like conformations, notably through phosphorylation by GSK3β^10^. In that study, phosphorylation of 2N4R tau promoted formation of fibrils with PHF-like morphological features, supporting a role for kinase activity in tau strain selection. However, GSK3β phosphorylates the proline-rich region less selectively than ERK2 and modifies a partially distinct set of sites, with more substantial phosphorylation also occurring at the PHF1 epitope in the C-terminal domain (Fig. S3). Consistent with this broader modification pattern, GSK3β-phosphorylated tau fibrils exhibited greater structural heterogeneity, precluding the determination of a single fibril fold at high resolution. In contrast, our findings demonstrate that a narrowly defined, kinase-specific phosphorylation pattern confined predominantly to the proline-rich region and excluding the amyloid core is sufficient to reproducibly encode a single dominant PHF fold. Together, these studies indicate that kinase identity, site selectivity, and regional targeting determine whether phosphorylation acts permissively or instructively in specifying tau fibril architecture.

Our results further distinguish native kinase-mediated phosphorylation from phosphomimetic and fragment-based models of tau aggregation. Short tau constructs can adopt PHF-like folds under specific conditions, but these systems lack the disordered flanking regions that constitute the fuzzy coat and influence fibril interactions, uptake, and seeding behavior^33–35^. Structural studies using full-length tau bearing limited phosphomimetic substitutions at the AT8 or PHF1 epitopes were insufficient to produce AD-like filaments, whereas more extensive phosphomimetic patterns were required to generate PHF folds in 3R but not 4R tau^8,27^. In contrast, phosphorylation of full-length 0N3R tau by a single, physiologically relevant kinase is sufficient to encode a PHF fold without cofactors or artificial substitutions. Notably, many ERK2 phosphorylation sites identified here overlap with phosphosites detected in tau isolated from Alzheimer’s disease brain, supporting the relevance of the modification pattern^17^.

Functionally, ERK2-phosphorylated tau fibrils display robust seeding activity in a cellular tau biosensor assay, providing independent validation that the structurally defined fibril state is competent for templated aggregation in cells. Notably, the enhanced seeding observed in biosensor cells correlates with the adoption of the AD PHF fold by ERK2-phosphorylated 0N3R tau rather than with increased fibril yield. These findings indicate that the specific ERK2-driven phosphorylation pattern promotes selection of a pathogenic PHF conformer, thereby enhancing seeding efficiency, instead of simply accelerating overall fibril formation.

More broadly, these findings support a general framework in which post-translational modifications of intrinsically disordered proteins encode amyloid polymorphism by acting on dynamic, non-core regulatory regions. In this view, kinase activity functions as a molecular gatekeeper that restricts access to particular folding trajectories, linking cellular signaling pathways to specific amyloid architectures without requiring direct chemical modification of the amyloid core.

An important open question raised by this work is whether kinase specificity is sufficient, in general, to encode alternative tau fibril folds. Tau is modified by multiple kinases *in vivo*, raising the possibility that combinations of phosphorylation events may act cooperatively or competitively to bias fold selection. Dissecting how individual and combinatorial kinase activities, potentially in combination with other post-translational modifications, shape tau fibril structure will be essential for defining the molecular basis of tau conformational diversity.

In summary, this work establishes that kinase-specific phosphorylation confined to a dynamic, non-core region of tau can encode selection of a defined amyloid fold. By providing a direct structural link between post-translational modification, fibril architecture, and seeding competence, our results advance a mechanistic understanding of tau strain formation and highlight phosphorylation as an instructive determinant of tau conformational diversity.

## Acknowledgments

We thank the Mass Spectrometry Facility of the Max Planck Institute for Multidisciplinary Sciences (MPI-NAT, Göttingen), in particular Dr. Olexandr Dybkov, for assistance with mass spectrometry analysis. We also thank Kerstin Overkamp from the Department of NMR-based Structural Biology, MPI-NAT Göttingen, for HPLC purification and intact mass spectrometry, the EM Facility of MPI-NAT for electron micrographs and the Facility for Light Microscopy at MPI-NAT Faßberg Campus. M.Z. was supported by the European Research Council (ERC) under the EU Horizon 2020 research and innovation program (Grant Agreement No. 787679). M.L. was supported by the Cure Alzheimer’s Fund, and the European Union (ERC, FuncAmyloid, 101087140). Views and opinions expressed are however those of the author(s) only and do not necessarily reflect those of the European Union or the European Research Council. Neither the European Union nor the granting authority can be held responsible for them. M.Z. and M.L. acknowledge research support from the Forschungskooperation Niedersachsen – Israel, Volkswagenstiftung, No: 76251-4659/2022 (ZN 4042). M.Z. acknowledges access to the 1.2 GHz spectrometer through the DFG Major Instrumentation Grant INST 1525/26-1 FUGG (project number 600373).

## Materials and Methods

### Protein Expression

Unlabeled human 0N3R tau was expressed in *E. coli* BL21™ (DE3) cells transformed with the pNG2 vector (a derivative of pET-3a, Merck-Novagen, Darmstadt) encoding the 0N3R tau isoform. Single colonies were used to inoculate LB medium supplemented with 100 μg/mL ampicillin and grown overnight at 37 °C. The overnight culture (22 mL) was transferred into 1 L of fresh LB medium and grown at 37 °C to an OD₆₀₀ of 0.8–0.9. Protein expression was induced with 1 mM IPTG for 1 hour at 37 °C. Cells were harvested by centrifugation, and pellets were stored at −80 °C until purification.

Uniformly ^15^N-labelled 0N3R tau was expressed using a similar protocol with a medium switch. After reaching an OD₆₀₀ of 0.8–0.9, cells were pelleted, washed with 1X M9 salts, and resuspended in M9 minimal medium containing ^15^NH₄Cl (1 g/L) as the sole nitrogen source. Following a 1-hour incubation at 37 °C, protein expression was induced with 1 mM IPTG and allowed to express overnight at 37 °C. Cells were then harvested and stored at −80 °C as described above.

Unlabelled 0N3R tau protein was purified as follows. Cell pellets were resuspended in lysis buffer containing 20 mM MES (pH 6.8), 1 mM EGTA, 2 mM DTT, 0.2 mM MgCl₂, protease inhibitor cocktail, lysozyme, and DNase I. Cells were lysed using a French press under ice-cold conditions. NaCl was added to the lysate to a final concentration of 500 mM, followed by heat treatment at 98 °C for 20 min to denature heat-labile proteins. Insoluble material was removed by ultracentrifugation at 127,000 g for 30 min at 4 °C.

The supernatant was treated with streptomycin sulfate (20 mg/mL) to precipitate nucleic acids, incubated for 15 min at 4 °C, and centrifuged at 14,000 g for 30 min. Tau protein was precipitated by the addition of ammonium sulfate (0.361 g/mL), incubated for 15 min, and pelleted by centrifugation at 14,000 g for 30 min. The pellet was resuspended in 20 mM MES (pH 6.8), 1 mM EDTA, 2 mM DTT, and 0.1 mM PMSF, and dialysed overnight against the same buffer. Dialysed samples were filtered and applied to a cation-exchange column (Mono S 10/100 GL, GE Healthcare) equilibrated in buffer A (20 mM MES pH 6.8, 1 mM EDTA, 2 mM DTT, 0.1 mM PMSF, 50 mM NaCl). Tau protein was eluted using a linear gradient up to 60% buffer B (20 mM MES pH 6.8, 1 M NaCl, 1 mM EDTA, 2 mM DTT, 0.1 mM PMSF). Tau-containing fractions were pooled, and further purification was performed by reverse-phase HPLC using a preparative C4 column (Vydac 214 TP, 5 μm, 8 × 250 mm) coupled to an ESI mass spectrometer. Protein purity was confirmed by mass spectrometry. Purified protein was lyophilised, resuspended in appropriate buffers, and stored at −80 °C. ^15^N-labelled 0N3R tau was purified using the same protocol up to cation-exchange chromatography. An additional size-exclusion chromatography step was performed using a HiLoad 26/200 Superdex 75 pg column equilibrated in PBS containing 500 mM NaCl. The tau-containing fraction was pooled and buffer-exchanged into 50 mM sodium phosphate containing 10 mM NaCl, and then stored at −80 °C.

### Phosphorylation of tau

Phosphorylation of 0N3R tau by ERK2 and GSK3β was performed using previously published protocols^17,36^, with minor modifications. Reaction conditions were optimised to favour site-specific phosphorylation while limiting hyperphosphorylation. For ERK2 phosphorylation, tau (200 µM) was incubated with ERK2 (0.017 mg mL⁻¹; E1283, Sigma-Aldrich or 14-550-M, Millipore) in buffer containing 50 mM HEPES (pH 8.0), 12.5 mM MgCl₂, 50 mM NaCl, and 2 mM DTT, supplemented with 2.5 mM ATP, 2 mM EGTA, and 1 mM PMSF. Reactions were carried out at 37 °C for 3 hours with shaking.

For GSK3β phosphorylation, tau (200 µM) was incubated with GSK3β (0.02 mg mL⁻¹; ab60863, Abcam) in buffer containing 25 mM HEPES (pH 7.2), 10 mM KCl, 5 mM MgCl₂, and 2 mM DTT, supplemented with 2 mM ATP, 5 mM EGTA, and 1 mM PMSF. Reactions were performed at 30 °C for 16 hours with shaking. Following phosphorylation, the kinase was inactivated by boiling the sample at 98 °C for 20 minutes. The precipitated protein was removed by centrifugation at 20,000 g for 30 minutes using an Eppendorf 5424 centrifuge. The resulting pellet was discarded, and the phosphorylated tau was buffer exchanged into the desired buffer using a Zeba Protein Desalting Column (Thermo Fisher).

### Aggregation assay

Unmodified and ERK2-phosphorylated tau samples were aggregated using a previously described cofactor-free protocol^18^. Tau protein (25 µM) was incubated at 37 °C in aggregation buffer containing 25 mM HEPES (pH 7.2), 10 mM KCl, 5 mM MgCl₂, 3 mM TCEP, and 0.01% NaN₃ in 96-well plates. Fibrillization was promoted by double-orbital shaking in the presence of three PTFE beads per well using a Tecan Spark plate reader. Thioflavin-T (50 µM) was added to monitor aggregation kinetics. All experiments were performed in triplicate.

### Fibril pelleting assay

Following the aggregation assay, the fibrils were pelleted by centrifugation at 20,000 g for 30 minutes using an Eppendorf centrifuge 5424. The concentration of soluble protein in the supernatant was determined using a micro BCA assay kit (Cat. No. 23235, Thermo Fisher Scientific). The amount of tau aggregated was calculated as the difference between the initial total tau concentration used in the aggregation assay and the soluble protein concentration found in the supernatant after fibril pelleting.

### Protease digestion of tau fibrils

Protease digestion of tau fibrils was performed following previously established procedures^18^. Briefly, 50 µl of 0.8 mg/mL fibril solution comprising unmodified or ERK2-phosphorylated 0N3R was supplemented with pronase (53702, Merck-Millipore) to a final concentration of 0.5 mg/mL. The digestion was carried out by incubating the solution at 37 °C for 30 minutes with shaking at 1440 rpm in an Eppendorf ThermoMixer. After the reaction, the protease activity was quenched by the addition of 1X cOmplete ™, EDTA-free Protease Inhibitor Cocktail. The protease-resistant fibril core was isolated by ultracentrifugation at 160,000 g for 30 minutes at 4 °C. The supernatant was carefully discarded, and the pellet was redissolved in aggregation assay buffer and loaded in a 15% SDS-PAGE gel. Later, the band containing the fibril core was excised from the SDS-PAGE gel and subjected to in-gel trypsin digestion before being loaded into the ESI mass spectrometer (Orbitrap Fusion Tribrid, Thermo Fisher) for peptide detection to identify the residues involved in the fibril core.

### HEK- biosensor cell seeding assay

A biosensor HEK293T cell line^24^ (ATCC® CRL-3275) stably expressing both tau RD P301S-CFP and tau RD P301S-YFP (amino acids: 244–372) was used to assess the seeding efficiency of ERK2-phosphorylated 0N3R tau fibrils. The ERK2-phosphorylated 0N3R fibrils were prepared as described earlier. Biosensor cells were seeded onto a poly-D-lysine-coated 24-well plate and maintained under standard mammalian cell culture conditions (37 °C, 5% CO2) for 24 hours. Prior to transfection, the fibrils were sonicated in a water bath (SONOREX DIGITECDT 102 H, Bandelin) for 2 minutes. The fibril seeds and control monomer were then mixed with Lipofectamine 2000 and incubated for 10 minutes. The resulting transfection mixture, containing 5 µg of fibrils/monomers, was added to each well. After 24 hours of incubation, fluorescence resonance energy transfer (FRET) images were acquired using a confocal microscope with excitation at 405 nm (CFP) and emission detection in the YFP range (520–550 nm). Image analysis and quantification were performed using Fiji (ImageJ).

### Mass Spectrometry

SDS-PAGE-separated protein samples were processed as described by Shevchenko et al. (1996)^37^. The resulting peptides were loaded onto nano HPLC (Dionex Ultimate 3000 UHPLC Thermo Fischer Scientific) coupled with an Orbitrap Mass spectrometer (Thermo Fisher). The peptides were separated with a linear gradient of 5–95% buffer B (80% acetonitrile and 0.08% formic acid) at a flow rate of 300 nL/min over a total gradient time of 58 min. To identify fibril core peptides, data analysis and searches were performed using Mascot (version 2.3.02), and the results were loaded into Scaffold (version 5.3.3) with 0N3R sequence.

The kinase-specific phosphorylation pattern of ERK2 in 0N3R using mass spectrometry was identified using the search engine PEAKS with a parent mass error tolerance of 10.0 ppm and a fragment mass error tolerance of 0.022 Da. Only peptides with a minimum Ascore of 20 or higher were considered for analysis to ensure accurate localisation of the phosphorylation site within the peptides. Residues consistently identified as phosphorylated across 3 independent mass spectrometry trials were confidently assigned as phosphorylation sites.

### NMR Spectroscopy

The 2D ^1^H–^15^N HSQC spectra of ^15^N-labelled unmodified and phosphorylated 0N3R tau (50 µM) were recorded in 50 mM sodium phosphate, 10 mM NaCl, and 1 mM TCEP (pH 6.8) at 298 K. Spectra of ERK2-phosphorylated 0N3R tau were acquired on a Bruker 1200 MHz spectrometer equipped with a 3 mm TCI cryoprobe, using 72 scans and acquisition times of td1 = 131.5 ms and td2 = 77.8 ms. Spectra of GSK3β-phosphorylated 0N3R tau were acquired on a Bruker 700 MHz spectrometer equipped with a 5 mm TCI cryoprobe, using 72 scans and acquisition times of td1 = 103 ms and td2 = 121.6 ms. Chemical shift assignments were based on previously published assignments for 2N4R tau and transferred to 0N3R tau^13^. Assignment transfer was validated by the consistency of chemical shifts. Residue-specific intensity ratios were calculated according to intensity ratio = 1− (I/I0), where I is the intensity of cross-peaks in the 2D 1H-15N HSQC spectrum of ERK2-phosphorylated tau and I0 is the intensity of the cross-peaks of unmodified tau. Spectral processing and analysis were performed using TopSpin 4.5.0 and POKY.

### Negative-stain Transmission Electron Microscopy (TEM)

The fibrils were sonicated for 2 minutes in an ultrasonic bath (SONOREX DIGITECDT 102 H, Bandelin), and 5 µL of 25 µM tau fibrils were treated with 1 µL of 1mg/mL pronase (Millipore) for 10 seconds before being applied to glow-discharged carbon-coated copper grids and stained with a 1% uranyl acetate solution. The EM images were captured using a F416 CMOS camera (TVIPS, Gauting, Germany) in conjunction with a CM 120 transmission electron microscope (FEI, Eindhoven, The Netherlands).

### Cryo-EM

The ERK2-phosphorylated 0N3R tau fibrils were concentrated four-fold by centrifugation of 100 µL of the fibril sample at 2500 g for 2 minutes and reducing the volume to 25 µL by removing the supernatant. The fibrils were further sonicated using a water bath (SONOREX DIGITECDT 102 H, Bandelin) for 2 minutes, and 5 µL of the sample was treated with 1 µL of 1 mg/mL pronase (Merck-Millipore) for 10 seconds. The mixture was then immediately added to glow-discharged Quantifoil R 2/1 200 mesh carbon copper grids. The grids were then plunge-frozen in liquid ethane using a Thermo Fisher Vitrobot. The cryo-EM grids were first screened to identify grids with well-dispersed fibrils using a Thermo Fisher Glacios TEM with a 200 kV field-emission gun. Final high-quality cryo-EM data were acquired using a Thermo Fisher Titan Krios G2 with a 300 kV field-emission gun. The images were captured with a slit width of 20 eV, a total dose of 40.0 electrons per Å², and dose fractionation of 40 frames (1.00 e/Å^2^/frame).

Recorded cryo-EM micrographs were processed in RELION using a standardised workflow^38,39^. Initial motion correction was carried out with RELION’s implementation of MotionCorr2, followed by estimation of contrast transfer function (CTF) parameters using CTFFIND4^40,41^. Fibrils were manually picked and extracted using a box size of 512 pixels and an inter-box distance of 13.6 pixels. Extracted particles were subjected to unbiased 3D classification with a reference-free cylinder to minimise model bias. Helical parameters were estimated by measuring fibril cross-over distances in the raw micrographs by applying a standard amyloid rise of 4.75 Å. From the resulting 3D classes, class averages containing continuous Cα atoms were picked, and 3D refinement was done using the 3D density map generated from the 3D classification. The helical twist and rise parameters were further refined for the resultant 3D refined density map with RELION’s search functions. The 3D refined density maps underwent Bayesian polishing and CTF refinement in RELION, followed by a final round of 3D map refinement. Final resolution estimates were determined using the 0.143 Fourier shell correlation criterion between half maps generated during 3D refinement, with a RELION-generated mask.

For atomic model building, an initial single-layer model of 0N3R tau fibril structure (PDB ID: 9CGX)^21^ was fitted to the cryo-EM reconstructed 3D density map. The model was refined in Coot (v0.9) and validated using comprehensive cryoEM tools in Phenix^42,43^. Iterative cycles of validation and adjustment in phenix and coot continued until Ramachandran and rotamer outliers, as well as structural clashes, were resolved. The validated model was replicated to generate a three-layer structure using Situs, with layers renamed using VMD^44,45^. Additional refinement and validation cycles were conducted in Phenix and Coot, and the middle layer was also adjusted in certain cycles with the ISOLDE plugin in ChimeraX to improve the model quality^46,47^. Iterative rounds of layer multiplication, validation, and adjustment were performed until a final model with zero Ramachandran, clash, and rotamer outliers and a low MolProbity score was obtained. All visualisation of the 3D density map and molecular model was performed using chimera and chimeraX^47,48^.

## Notes

https://www.rcsb.org/structure/unreleased/31PS

### Competing Interest Statement

The authors have declared no competing interest.

